# BBSome regulation of LITE-1 receptor in a cilium-independent manner in *C. elegans*

**DOI:** 10.1101/2020.04.27.064998

**Authors:** Xinxing Zhang, Jinzhi Liu, Jianfeng Liu, X.Z. Shawn Xu

**Author notes:** These authors contributed equally to this paper (Xinxing Zhang, Jinzhi Liu).

## Abstract

Bardet-Biedl Syndrome (BBS) is a genetic disorder affecting primary cilia. BBSome, a protein complex composed of eight BBS proteins, regulates the structure and function of cilia in diverse organisms, and its malfunction causes BBS in humans. Here, we report a new function of BBSome in *C. elegans*. In a forward genetic screen for genes regulating the light sensitivity of the ciliated ASH sensory neurons, we isolated *bbs* mutants, indicating that BBSome regulates ASH photosensitivity. Surprisingly, cilia are not required for ASH neurons to sense light, suggesting that BBSome regulates ASH photosensitivity independently of cilia. Interestingly, the light-sensing receptor LITE-1, which mediates photosensation, is a non-ciliary protein in ASH neurons. LITE-1 in ASH neurons becomes unstable in *bbs* mutants in an age-dependent manner, indicating that BBSome regulates the stability of LITE-1 in these neurons. These results identify a cilium-independent function of BBSome in regulating a non-ciliary protein in ciliated cells.

## Introduction

Bardet-Biedl Syndrome (BBS) is an autosomal recessive genetic disorder characterized by a wide spectrum of clinical symptoms. Retinal degeneration, obesity, polydactyly (extra fingers or toes), hypogonadism, kidney malfunction, and learning disabilities are considered the primary symptoms of BBS ^1^. At least twenty-one genes are associated with BBS ^2^. Eight BBS proteins form a stable core complex termed BBSome ^3,4^. BBSome is enriched in the basal body of cilium and centrosome and is important for maintaining the function of cilia, and to a lesser extent the structure of cilia ^3–6^. For example, BBSome facilitates the transport of cargo proteins into cilia and is also required for retrieving signaling molecules from cilia ^5,7–12^. BBSome has also been shown to be associated with intraflagellar transport (IFT), and undergoes bidirectional movement along cilia in mammalian neurons ^13^. Thus, BBSome maintains the proper function of cilia mainly by regulating protein trafficking both in and out of cilia. Nevertheless, the pleiotropic symptoms of BBS patients raise the possibility that some of the symptoms might result from non-ciliary functions of BBSome.

The nematode *C. elegans* represents a highly valuable genetic model organism for the study of BBS ^14,15^. Eight highly conserved *bbs* genes are found in the *C. elegans* genome ^14,15^. Among them, seven encode BBSome components, and the other one encodes a homolog of the human GTPase BBS3/Arl6, ARL-6 ^14,15^. All *bbs* genes are exclusively expressed in ciliated neurons under the control of the transcription factor DAF-19 ^6,16,17^. BBSome is enriched in the base of cilia and is also localized along the axoneme of cilia ^16,18^. Disruption of BBSome in worms results in slightly truncated cilia ^19^. Similar to mammalian BBSome, worm BBSome directly interacts with IFT sub-complexes, coordinates IFT complex movement in the cilia, and is essential for IFT complex assembly ^18–20^. Mutations in worm *bbs* genes cause pleotropic phenotypes, some of which are reminiscent of those observed in BBS human patients, such as obesity and learning disabilities ^21–23^. BBSome mutant worms also display moderate sensory defects at the behavioral level ^19,24^. Work in *C. elegans* also shows that *bbs* genes regulate dense-core vesicle secretion in ciliated neurons independently of the loss of cilia via an unknown mechanism, pointing to a non-ciliary function of *bbs* genes ^21^. Nevertheless, the precise role of *C. elegans* BBSome in regulating cilium structure and function is incompletely understood, and it is also unclear to what extent and how BBSome mediates non-ciliary functions in ciliated neurons.

Here, we report a new function of BBSome in *C. elegans*. In a genetic screen for genes regulating the light sensitivity of the ciliated ASH sensory neurons, we isolated several *bbs* mutants. Further analysis shows that mutations in each components of BBSome all result in a defect in ASH light sensitivity, but not other sensory functions of ASH neurons. Notably, ASH neurons do not require cilia to sense light, suggesting that *bbs* genes regulate ASH light sensitivity in a cilium-independent manner. In support of this model, we find that the photoreceptor protein LITE-1, which is the primary light sensor in *C. elegans* ^25^, is a non-ciliary protein in ASH neurons, suggesting that LITE-1 mediates light sensation independently of cilia in these ciliated neurons. In *bbs* mutants, ASH neurons progressively lose LITE-1 protein in an age-dependent manner, indicating that BBSome is required for maintaining the stability of LITE-1. Our results reveal an unexpected function of BBSome in regulating the stability of non-ciliary proteins in ciliated cells. This supports the notion that some of the pleotropic symptoms of BBS may result from non-ciliary functions of BBSome.

## Results

### BBSome is required for ASH neurons to sense light

Despite the lack of eyes, worms are sensitive to short-wavelength light and engage in negative phototaxis behavior to avoid light ^26^. Worms sense light through the photoreceptor protein LITE-1, a unique type of light sensor ^25,27,28^. At least four pairs of ciliated sensory neurons, ASH, AWB, ASK and ASJ, are required for light-evoked avoidance response in the head region ^26^. Among them, ASH is a bit special in that it is a polymodal nociceptive neuron, sensing nearly all types of aversive cues, including odorants, tastants, mechanical forces, high osmolarity, pH, etc ^29,30^. While the photosensation mechanisms in some photoreceptor neurons such as ASK and ASJ have been well characterized ^28^, little is known about how ASH neurons sense light. To address this question, we first confirmed that ASH neurons are indeed light-sensitive, as light evoked robust calcium response in these neurons (Figure 1A-B). ASH photosensitivity was not altered in *unc-13* and *unc-31* mutants, which is defective in secreting small neurotransmitters and neuropeptides, respectively ^31,32^, indicating that the observed light response in ASH neurons primarily arose from ASH neurons cell-autonomously (Figure 1A-B). As expected, LITE-1 is required for ASH neurons to sense light, as *lite-1(xu492)* mutant, a deletion allele of *lite-1*, did not respond to light stimulation (Figure 1A-B).

**Figure 1.**
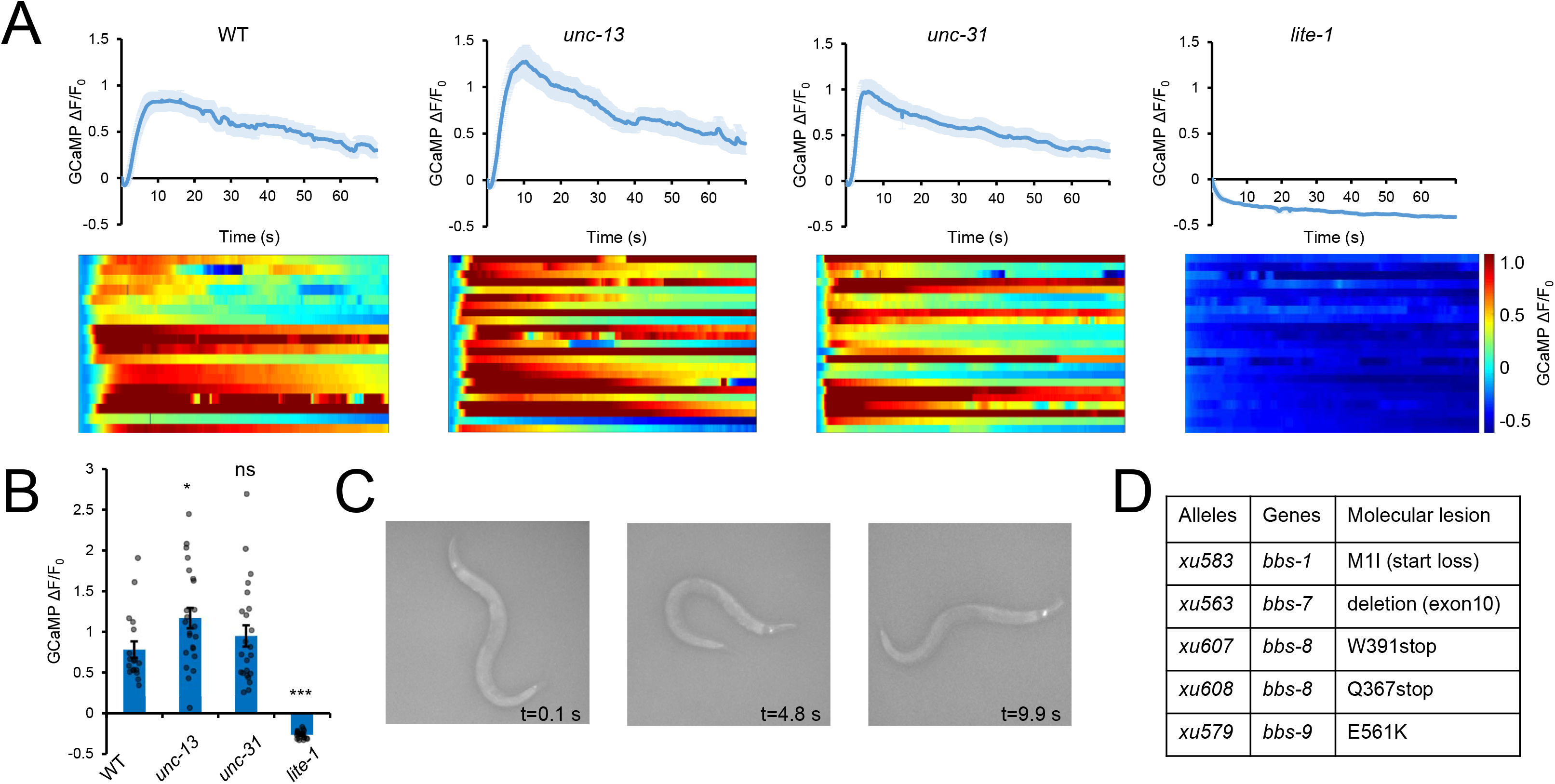
*bbs* genes are required for ASH neurons to sense light. **(A)** ASH neurons sense light cell-autonomously in a LITE-1-dependent manner. Calcium imaging was performed on wild type, *unc-13(e51)*, *unc-31(e169),* and *lite-1(xu492)* worms. GCaMP6f was expressed as a transgene in ASH neurons using the *sra-6* promoter. Worms were imaged and stimulated with blue light. Top panels: averaged calcium traces. Shades along the traces denote error bars (SEM). Bottom panel: heat map showing individual traces from each tested animals. n≥18. **(B)** Bar graph summarizing the data in (A). Error bars: SEM. Statistics were calculated using One-way ANOVA Bonferroni test, and all strains were compared to wild type. ***P<0.0001, *P<0.05, ns: not significant. **(C)** Snapshot images. Upon blue light illumination, the fluorescence in ASH neurons in a moving worm turned brighter, enabling a genetic screen for mutants insensitive to light stimulation. The strain carried a transgene expressing the genetically-encoded calcium sensor Case12 in ASH neurons under the *sra-6* promoter. **(D)** Five mutant alleles identified from the genetic screen were mapped to *bbs* genes, including *bbs-1(xu583), bbs-7(xu563), bbs-8(xu607), bbs-8(xu608)*, and *bbs-9(xu579).*

We then designed a forward genetic screen to identify genes that regulate ASH photosensitivity. We labeled ASH neurons with a GECI (genetically-encoded calcium indicator) that emits fluorescence excited by blue light (Figure 1C). As ASH neurons show sensitivity to blue light, we were able to detect an increase in fluorescence intensity in ASH neurons upon light stimulation (Figure 1C). Taking advantage of this observation, we mutagenized worms with EMS and screened for mutants that failed to exhibit light-evoked increase in fluorescence intensity. After going through ~36,000 F2 worms (from ~2000 F1), we isolated 14 mutants. Among them, two (i.e. *xu580[S320F]* and *xu592[P112L]*) are *lite-1* mutants, validating the screen. Interestingly, we isolated five mutants that were mapped to *bbs* genes, including *bbs-1, bbs-7, bbs-8,* and *bbs-9* (Figure 1D). Since we hit half of the genes that encode protein products of *C. elegans* BBSome, we wondered if all the *bbs* genes are required for ASH neurons to sense light. Remarkably, we found that seven out of eight BBSome genes are essential for ASH neurons to respond to light stimulation (Figure 2). Loss of *arl-6*, the remaining BBSome-associated gene that encodes a homolog protein of human GTPase BBS3/Arl6, also led to a defect in ASH photosensitivity (Figure 2C and 2I). Thus, it appears that all BBSome components are important for ASH neurons to sense light.

**Figure 2.**
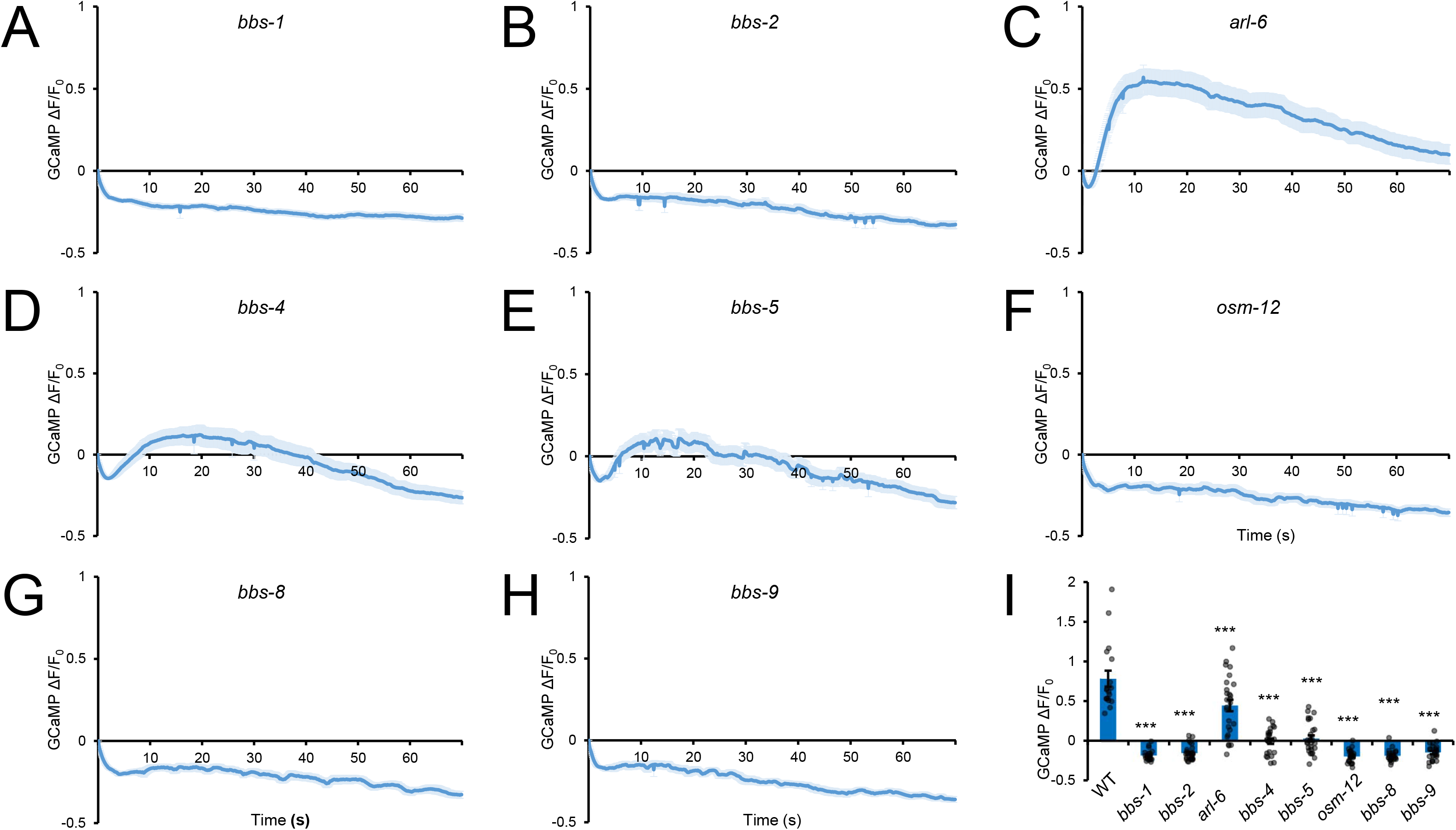
BBSome is required for ASH neurons to sense light. **(A-H)** Calcium imaging of *bbs-1(ok1111)*, *bbs-2(ok2053)*, *arl-6/bbs-3(ok3472)*, *bbs-4(tm3038)*, *bbs-5(gk537)*, *osm-12/bbs-7(ok1351)*, *bbs-8(nx77)*, and *bbs-9(gk471)* mutant worms. All strains carried the same GCaMP transgene. Shown are calcium imaging traces. Shades along each trace denote error bars (SEM). **(I)** Bar graph summarizing the data in (A-H). Error bars: SEM. n≥18. Statistics were calculated using One-way ANOVA Bonferroni test, and all strains were compared to wild type. ***P<0.0001, **P<0.005, *P<0.05.

### A specific role of BBSome in regulating light sensation in ASH neurons

Since ASH neurons are polymodal sensory neurons, we asked whether BBSome is important for ASH neurons to sense other sensory cues, such as high osmolarity (glycerol), heavy metals (copper), and detergents (SDS). We focused on BBS-1 and BBS-8, two core components of BBSome. Both *bbs-1* and *bbs-8* mutants responded robustly to all tested stimuli (Figure 3). As a control, *osm-9* mutant worms exhibited a severe defect (Figure 3). We also found that though ASH neurons lost light sensitivity in *bbs* mutants, ASK neurons responded normally to light in these mutants (Figure S1A-B). These data together reveal a specific role for BBSome in regulating light sensation in ASH neurons.

**Figure 3.**
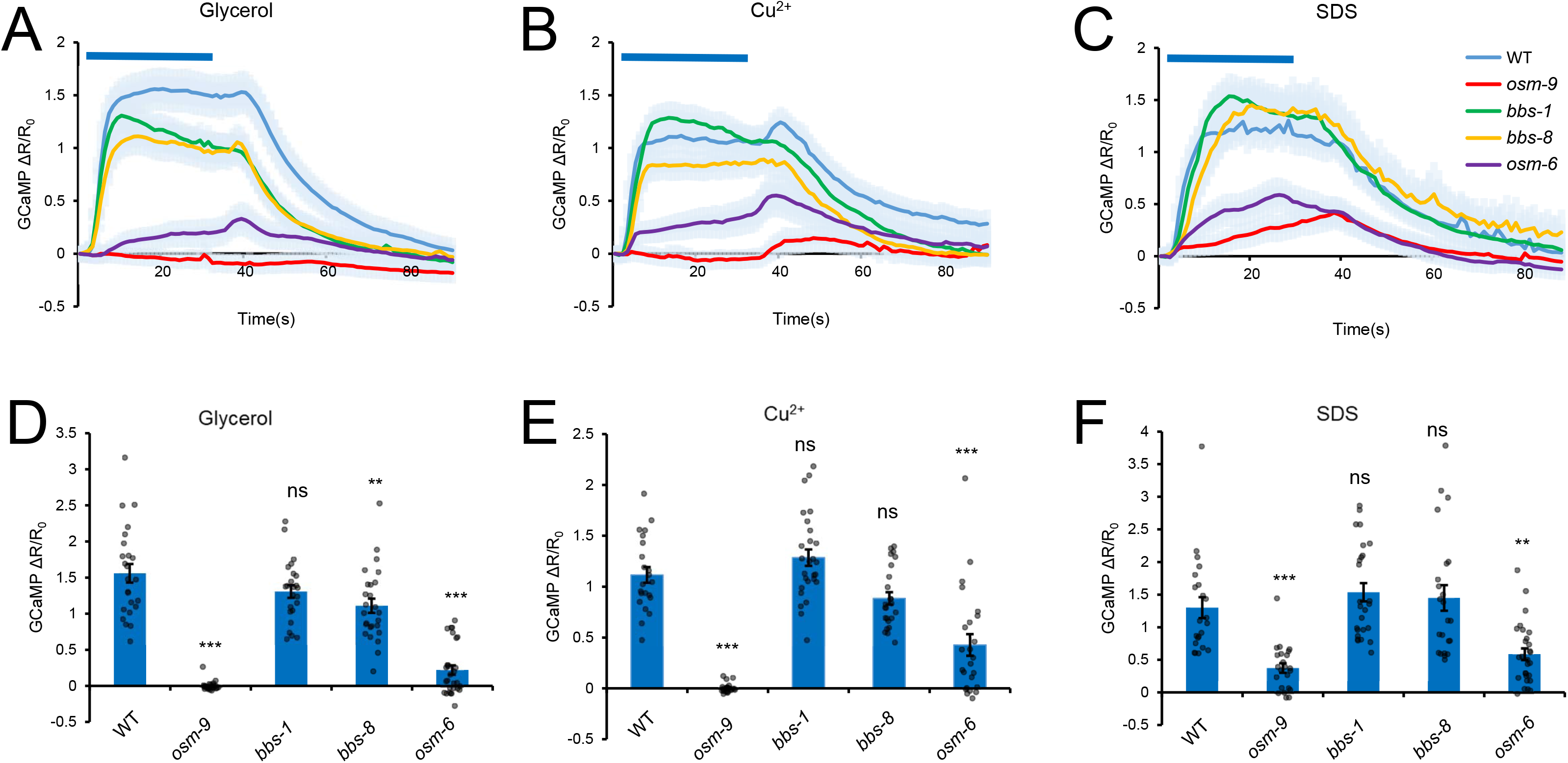
BBSome is not required for ASH neurons to sense aversive cues such as high osmolarity (glycerol), heavy metals (copper), and detergents (SDS). **(A-C)** ASH neurons in *bbs-1* and *bbs-8* mutant worms respond normally to glycerol (A), copper (B) and SDS (C). All strains carried the same GCaMP transgene. Worms were pre-exposed to blue light to quench the light response to establish a basal line. Subsequently, worms were stimulated by 1 M glycerol, 1 mM Cu^2+^ or 0.1% SDS. *osm-9* and *osm-6* worms were included as controls. Shown are average traces. Shades along the traces denote error bars (SEM). **(D-F)** Bar graphs summarizing the data in (A-C). Error bars: SEM. Statistics were calculated using One-way ANOVA Bonferroni test, and all strains were compared to wild type. n≥22. *** indicates P<0.0001, ** indicates P<0.005, * indicates P<0.05, and ns indicates not significant.

### Cilia are not required for ASH to sense light

In *C. elegans*, BBSome is required for IFT (intraflagellar transport), which is important for building and maintaining the structure and function of cilia ^14,15^. The observation that *bbs* mutants lacked photosensitivity in ASH neurons suggests that cilia are essential for ASH neurons to sense light. To test this idea, we examined cilium-defective mutants, particularly those affecting IFT, such as *osm-3* (kinesin motor), *che-3* (dynein motor), *daf-10* (IFT-A component), *osm-5* (IFT-B component), *klp-11* (kinesin-II motor), and *kap-1* (kinesin-associated protein) (Figure 4A-B). In our EMS genetic screens, we conducted a counter screen to exclude mutants with severe defects in ASH morphology, which may explain the fact that we did not recover these cilium-defective mutants. Here, we found that while mutations in *osm-3, che-3, daf-10* and *osm-5* rendered ASH neurons insensitive to light, *klp-11* and *kap-1* mutant worms showed normal response to light (Figure 4A-B). Thus, while some IFT components (e.g. BBSome, OSM-3, and DAF-10) are important for ASH neurons to sense light, others (KLP-11 and KAP-1) are not, suggesting that ASH photosensitivity does not necessarily require every IFT component.

**Figure 4.**
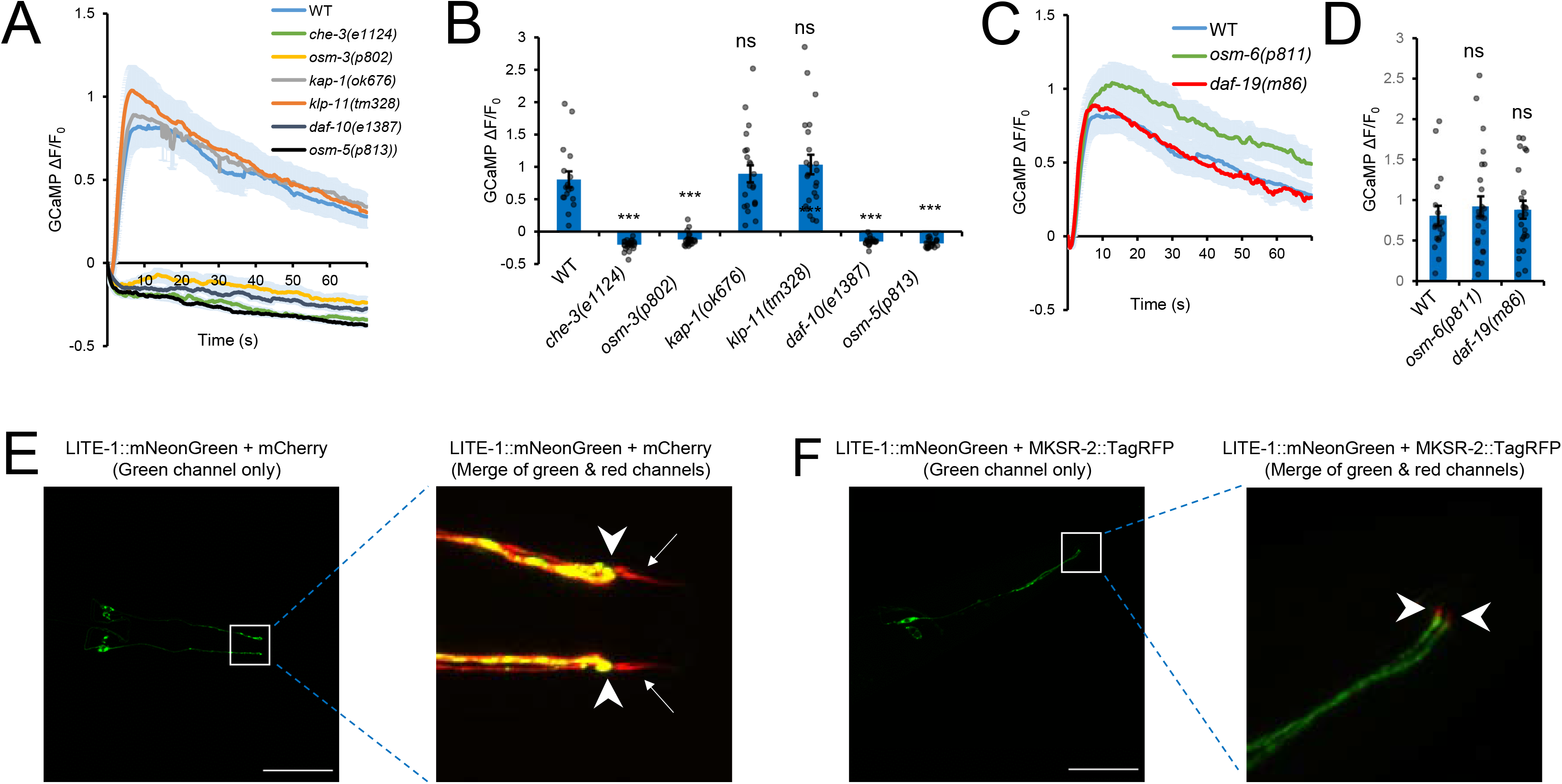
Cilia are not required for ASH neurons to sense light and LITE-1 is a non-ciliary protein in ASH neurons. **(A-B)** Not all IFT mutants are defective in light sensation in ASH neurons. All strains carried the same GCaMP transgene. (A) Calcium imaging traces with shades along the traces denoting error bars (SEM). (B) Bar graph. Error bars: SEM. n≥17. Statistics were calculated using One-way ANOVA Bonferroni test, and all strains were compared to wild type. ***P<0.0001. ns: not significant. **(C-D)** *osm-6(p811)* and *daf-19(m86)* mutant worms respond normally to light in ASH neurons. All worms carried the same GCaMP transgene. (C) Calcium imaging traces. The shades along the traces denote error bars (SEM). (D) Bar graph. Error bars: SEM. n≥17. **(E)** LITE-1 is not localized to ASH cilia. Shown are confocal images of transgenic worms, in which LITE-1∷mNeonGreen fusion protein was expressed in ASH neurons together with mCherry under the *sra-6* promoter. Shown on the right is a zoomed-in image. Arrows point to the cilia where LITE-1∷mNeonGreen signals (green) are absent. Arrowheads point to the junction between the dendrite and cilia. Scale bar: 50 μm. **(F)** LITE-1 is not localized to the transition zone in ASH. Shown are confocal images of transgenic worms, in which LITE-1∷mNeonGreen fusion protein was expressed in ASH neurons together with MKSR-2∷TagRFP fusion protein under the *sra-6* promoter. Shown on the right is a zoomed-in image. MKSR-2 is a transition zone marker. Arrow heads point to the transition zone (red) where LITE-1∷mNeonGreen (green) signals are absent. Scale bar: 50 μm.

To further evaluate the role of cilia in ASH photosensitivity, we tested OSM-6, a core component of the IFT-B subcomplex ^33^. *osm-6* mutants are commonly used for assessing ciliary functions, as their cilia are nearly eliminated (much shortened) ^33,34^. Intact ciliary structure is believed to be required for ASH neurons to sense sensory cues. In support of this notion, *osm-6(p811)* mutant worms were defective in responding to a number of aversive sensory cues, such as high osmolarity (glycerol), heavy medals (copper), and detergents (SDS) (Figure 3). Surprisingly, ASH neurons responded normally to light in *osm-6(p811)* mutant worms (Figure 4C-D), suggesting that intact cilia are not required for ASH neurons to sense light.

As *osm-6(p811)* mutant worms still retain some rudimentary ciliary structure ^34^, it might be argued that such defective cilia might nevertheless be sufficient to allow ASH neurons to sense light. To address this concern, we checked DAF-19, a transcription factor known to be required for the formation of cilia in all ciliated neurons. It has been reported that *daf-19(m86)* mutant worms fail to develop any type of ciliary structure, including transition zones ^34^. Remarkably, these mutant worms responded normally to light (Figure 4C-D). Thus, we conclude that cilia are not required for ASH neurons to sense light.

Another interesting observation concerns BBS-4 and BBS-5. Unlike other BBSome components, BBS-4 and BBS-5 are functionally redundant and thus not required for ciliogenesis, and both *bbs-4* and *bbs-5* mutants show normal ciliary structure ^35^. However, despite the presence of intact cilia in *bbs-4* and *bbs-5* mutant worms, ASH neurons in these two mutants failed to respond to light (Figure 2D, 2E, and 2I). Thus, cilia are neither necessary nor sufficient for ASH neurons to sense light.

### LITE-1 is a non-ciliary protein in ASH neurons

The fact that ASH neurons do not depend on cilia for light sensation raises the possibility that LITE-1 does not function in cilia. We thus checked the subcellular localization of LITE-1 protein in ASH neurons. We expressed LITE-1∷mNeonGreen fusion as a transgene in ASH neurons. mCherry was co-expressed as a marker to label the entire neuron. We found that LITE-1∷mNeonGreen fusion did not overlap with mCherry in the cilia (Figure 4E), indicating that that LITE-1 was not localized to the cilia in ASH neurons. To ascertain whether LITE-1 can enter the transition zone, we generated another transgenic strain, in which the transition zone was labeled by MKSR-2∷TagRFP fusion in ASH neurons (Figure 4F). LITE-1 did not appear to overlap with MKSR-2 (Figure 4F). These data demonstrate that LITE-1 is not localized to ASH cilia or transition zone, identifying LITE-1 as a non-ciliary protein in ASH neurons. This is consistent with the observation that cilia are not required for ASH neurons to sense light.

### Age-dependent loss of LITE-1 protein in ASH neurons in *bbs* mutants

Since LITE-1 is a non-ciliary protein in ASH neurons and cilia are neither necessary nor sufficient for ASH neurons to sense light, this brings up the question of how BBSome regulates LITE-1. To assess whether and how disruption of BBSome affects LITE-1 in ASH neurons, we fused an mNeonGreen tag to the C-terminus of the endogenous LITE-1 receptor by CRISPR/Cas9-based genome editing. The resulting strain, named *lite-1(xu708)*, showed normal phototaxis behavior, indicating that the mNeonGreen tag did not notably affect LITE-1 function (Figure S2). LITE-1∷mNeonGreen fluorescence was very dim, suggesting that endogenous LITE-1 is expressed at a very low level, consistent with previous work ^25^. Notably, ASH is among those few cells in which LITE-1 was expressed at a relatively higher level, making it possible to visually identify this neuron (Figure 5A). Again, we focused on characterizing BBS-1 and BBS-8, two core components of BBSome. At the larval stages up to L4, we detected LITE-1 protein in ASH neurons of *bbs* mutant and wild-type worms (Figure 5A). However, at the adult stage beginning at day 2, we can no longer detect LITE-1 protein in ASH neurons of *bbs* mutants, though LITE-1 expression in ASH neurons persisted in wild-type worms (Figure 5A). As an internal control, we found that LITE-1 expression was normal in ASK neurons of *bbs* mutants at both the larval and adult stages (Figure 5A). Thus, mutations in *bbs* genes cause an age-dependent loss of LITE-1 expression in ASH neurons.

**Figure 5.**
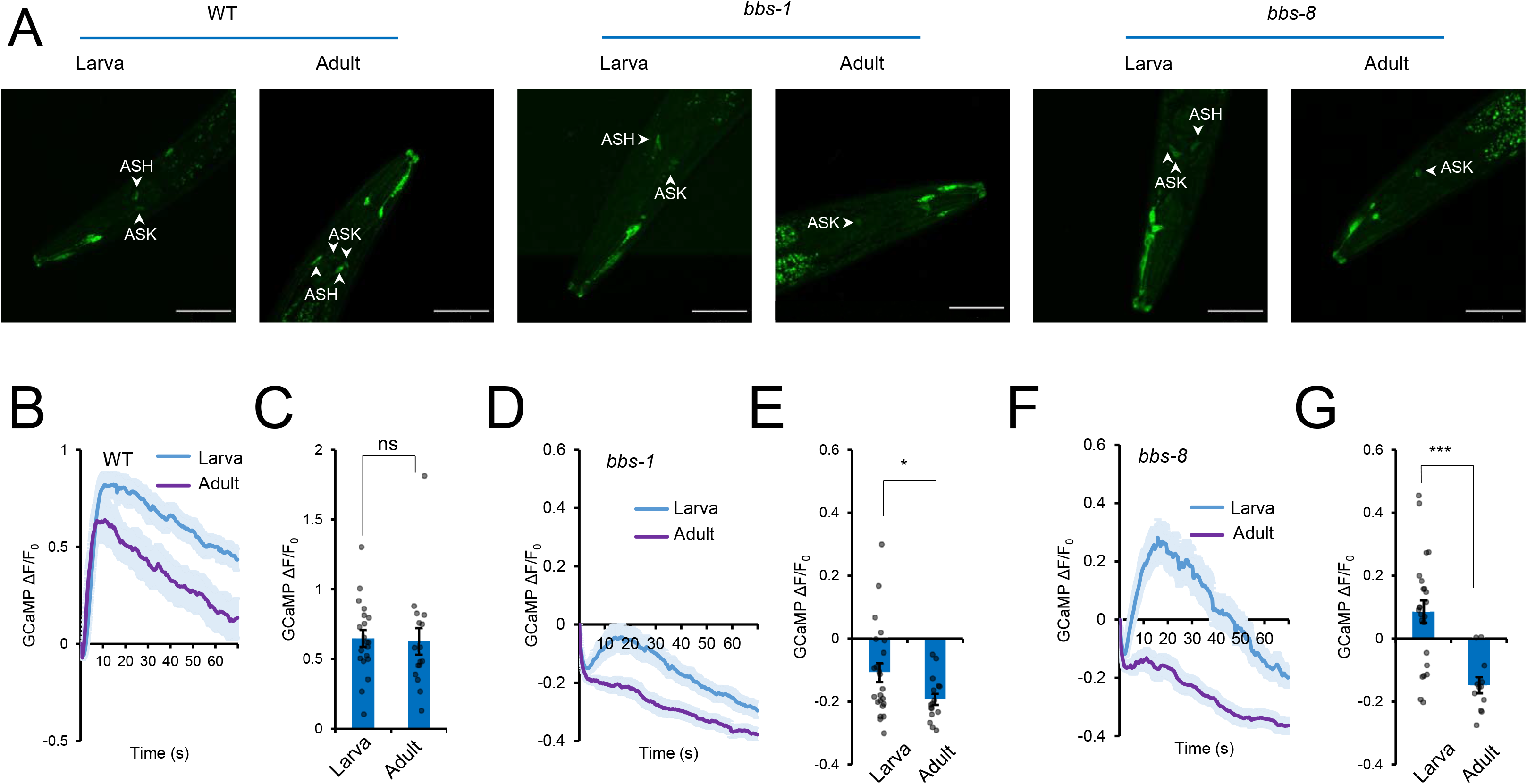
Progressive loss of LITE-1 protein in ASH neurons of *bbs* mutants in an age-dependent manner. **(A)** LITE-1 protein is present at the larval stage but absent at the adult stage in *bbs* mutant worms. Shown are confocal images for *lite-1(xu708[LITE-1∷mNeonGreen])* in wild-type, *bbs-1(ok1111)*, and *bbs-8(nx77)* genetic backgrounds at both the larval (L4) stage and adult (day 2) stage. *lite-1(xu708[LITE-1∷mNeonGreen])* is a CRISPR/Cas9 knock-in strain, in which an mNeonGreen tag is fused to the C-terminus of LITE-1. LITE-1 protein was found in both ASH and ASK neurons in wild-type worms at both the larval and adult stages. However, in *bbs* mutants, LITE-1 in ASH neurons can only be detected at the larval stage, but not at the adult stage. Scale bar: 50 μm. **(B-C)** ASH neurons in wild-type worms at both the larva and adult stages respond to light. (B) ASH calcium imaging traces from worms at the larva (L4) stage and adult (day 1) stages. Shades along the traces denote error bars (SEM). (C) Bar graph. Error bars: SEM. n≥16. **(D-E)** ASH neurons in *bbs-1* mutant worms at the larval stage respond to light, but those at the adult stage do not. (D) Calcium imaging traces from *bbs-1(ok1111)* mutant worms at the larval (L4) and adult (day1) stages. Shades along the traces denote error bars (SEM). (E) Bar graph. n≥16. Error bars: SEM. *P<0.05 (t test). **(F-G)** ASH neurons in *bbs-8* mutant worms at the larval stage respond to light, but those at the adult stage do not. (F) Calcium imaging traces from *bbs-8(nx77)* mutant worms at the larval (L4) and adult (day1) stages. Shades along the traces denote error bars (SEM). (G) Bar graph. n≥12. Error bars: SEM. ***P<0.0001 (t test).

We then assayed the *lite-1* mRNA level in *bbs* mutants. To do so, we first isolated ASH neurons by fluorescence-activated cell sorting (FACS) ^36^, and then extracted their RNA. qRT-PCR analysis detected no significant difference in the *lite-1* mRNA level between wild-type and *bbs* mutants in adult worms (Figure S3), indicating that *bbs* mutations do not affect the mRNA level of *lite-1*. This suggests that BBSome is important for maintaining the stability of LITE-1 protein in ASH neurons. Consistent with this notion, at the functional level, we found that ASH neurons indeed responded to light in *bbs* mutants at the larval stage, though no such light response was detected in adult worms of these mutants (Figure 5B-G). These results together unveil an important role for BBSome in regulating the stability of LITE-1 protein in ASH neurons.

If BBSome is important for maintaining LITE-1 stability in ASH neurons, then overexpression of LITE-1 should be able to bypass the requirement of BBSome in ASH neurons but not *vice versa*. To test this hypothesis, we generated a transgene driving LITE-1 expression in ASH neurons. As expected, this transgene rescued the light-sensing defect of ASH neurons in *lite-1* mutant worms (Figure 6A-B). Remarkably, this same *lite-1* transgene also rescued the ASH light-sensing defect in *bbs* mutants (Figure 6A-B). By contrast, overexpression of *bbs* genes in ASH neurons failed to rescue the ASH light-sensing defect observed in *lite-1* mutant worms, though these *bbs* transgenes were able to rescue the defect in *bbs* mutants (Figure 6A-B). Thus, overexpression of LITE-1 can bypass the requirement of BBSome in ASH neurons but not *vice versa*, suggesting that BBSome may act upstream of LITE-1 to regulate its stability.

**Figure 6.**
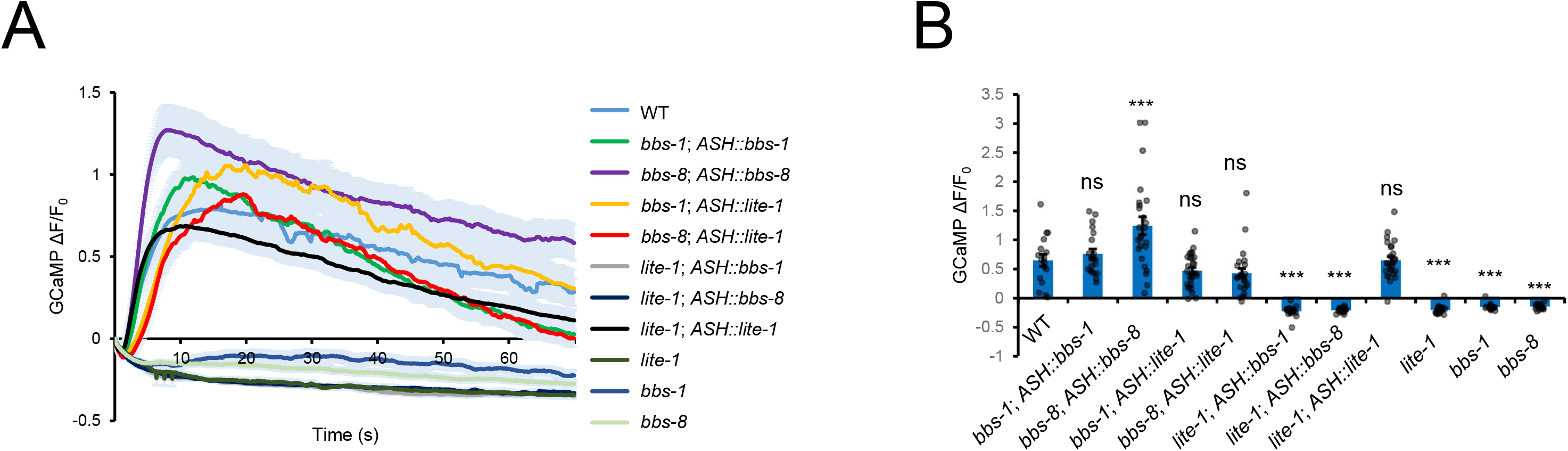
BBSome acts upstream of LITE-1 to regulate ASH light sensitivity. **(A)** Overexpression of LITE-1 in ASH can rescue the light insensitivity phenotype in *bbs* mutants, but not *vice versa*. The light-sensing defects in ASH neurons in both *bbs-1(ok1111)* and *bbs-8(nx77)* mutants can be rescued by expressing either *bbs-1* and *bbs-8* cDNA or by expressing *lite-1* cDNA. However, the light-sensing defects in ASH neurons in *lite-1* mutant worms cannot be rescued by *bbs-1* or *bbs-8* cDNA, though both cDNA can rescue the defect in *bbs* mutants. Shown are calcium imaging traces. Shades along the traces denote error bars (SEM). **(B)** Bar graph summarizing the data in (A). Error bars: SEM. Statistics were calculated using One-way ANOVA Bonferroni test, and all strains were compared to wild type. n≥17. ***P<0.0001; **P<0.005; *P<0.05; ns: not sign

## Discussion

BBSome is well known for its role in maintaining the function and structure of cilia. Given that BBS patients manifest a wide spectrum of clinical symptoms such as retinal degeneration, obesity, polydactyly, hypogonadism, kidney malfunction, and learning disabilities, it has been suggested that BBSome may also function outside of cilia ^1^. However, it is unclear whether, to what extent, and how non-ciliary functions of BBSome may contribute to such diverse symptoms of BBS patients. Here, we identified a cilium-independent function of BBSome in regulating the stability of a non-ciliary protein in ciliated cells in *C. elegans*. Specifically, we show that BBSome mutants exhibit an age-dependent loss of photosensitivity in the ciliated ASH sensory neurons. ASH neurons do not require cilia for light sensing, which is mediated by the photoreceptor protein LITE-1, a non-ciliary protein. We further show that in BBSome mutants, ASH neurons progressively lose LITE-1 protein but not *lite-1* mRNA in an age-dependent manner and hence their sensitivity to light, demonstrating that BBSome regulates LITE-1 stability. Interestingly, overexpression of LITE-1 can bypass the requirement of BBSome in ASH neurons but not *vice versa*, suggesting that BBSome acts upstream of LITE-1 to regulate its stability. These results together uncover a novel cilium-independent role of BBSome in maintaining the stability of a non-ciliary protein in ciliated cells.

One interesting observation is that BBSome appears to be specifically required for ASH neurons to sense light but not other types of aversive cues. Worms with disrupted BBSome have been reported to develop slightly truncated cilia revealed by some fluorescent protein markers ^19^. However, more recent work by transmission electron microscopy (TEM) demonstrates that in *bbs* mutants, the overall structure of cilia is very similar to that in wild-type worms, a phenomenon that is very different from mutants lacking some other IFT proteins ^37^. At the functional level, the molecular machinery (e.g. ODR-3 and OSM-9), which mediates ASH sensing of other aversive cues, is enriched in ASH cilia ^29^. Though malfunction of BBSome causes signaling protein mislocalization, such mislocalization is usually confined to the cilia, and the signaling proteins remain accumulated within the cilia, in which case they may still be able to function in sensory transduction ^38^. On the other hand, as LITE-1 is a non-ciliary protein and disruption of BBSome leads to a progressive loss of LITE-1 protein, *bbs* mutants would then become insensitive to light. This may provide a potential explanation to the observation that ASH neurons lose photosensitivity but respond normally to other aversive cues in *bbs* mutant worms. Another interesting observation is that while *bbs* genes are ubiquitously expressed in all ciliated neurons ^6,16,17^, they are required for ASH neurons, but not some other photoreceptor neurons such as ASK, to sense light. Since both BBSome and LITE-1 are expressed in these neurons, it is unclear how the specificity is achieved. It is possible that some unique regulators are present in ASH neurons to endow BBSome with the ability to regulate LITE-1 stability. Future studies are needed to elucidate the detailed mechanisms.

In humans, BBS patients also progressively lose vision ^1^. The patients usually begin to lose night vision at an age of ~8.5 years old and become blind when reaching ~15.5 years old ^1^. This phenomenon has also been observed in various mouse models of BBS. Retinal degeneration and signaling protein mislocalization are commonly associated with vision loss in these mouse models ^39–41^. Interestingly, photoreceptor cells display an age-dependent loss of the photoreceptor protein rhodopsin in a BBS mouse model ^41^, revealing an interesting analogy between worm and mouse BBSome in regulating the stable expression of photoreceptor proteins. Currently, this role of mammalian BBSome has been attributed to its function in the cilia. Our results raise the possibility that a cilium-independent function of BBSome may also contribute to the age-dependent vision loss observed in mouse BBS models and perhaps human patients.

## Materials and Methods

### Strains

TQ3030 N2
TQ9258 *bbs-1(ok1111)* I
TQ9255 *bbs-8(nx77)* V
TQ8029 *lite-1(xu492)* X
TQ9387 *lite-1*(*xu708*[LITE-1∷mNenoGreen∷3XFlag]) X
TQ8866 N2 *xuIs556*[*Psra-6∷Case12*+*Psra-6∷SL2∷mCherry*]
TQ8979 *bbs-1(xu583)* I; *xuIs556*[*Psra-6∷Case12*+*Psra-6∷SL2∷mCherry*]
TQ8947 *osm-12(xu563)* III; *xuIs556*[*Psra-6∷Case12*+*Psra-6∷SL2∷mCherry*]
TQ9028 *bbs-8(xu607)* V; *xuIs556*[*Psra-6∷Case12*+*Psra-6∷SL2∷mCherry*]
TQ9030 *bbs-8(xu608)* V; *xuIs556*[*Psra-6∷Case12*+*Psra-6∷SL2∷mCherry*]
TQ8972 bbs-9(xu579) I; *xuIs556*[*Psra-6∷Case12*+*Psra-6∷SL2∷mCherry*]
TQ5856 N2; *xuEx1978*[*Psra-6∷GCaMP6f*+*Psra-6∷SL2∷DsRed*]
TQ9613 *unc-13(e51)* I; *xuEx1978*[*Psra-6∷GCaMP6f*+*Psra-6∷SL2∷DsRed*]
TQ9614 *unc-31(e169)* IV; *xuEx1978*[*Psra-6∷GCaMP6f*+*Psra-6∷SL2∷DsRed*]
TQ8220 *lite-1(xu492)* X; *xuEx1978*[*Psra-6∷GCaMP6f*+*Psra-6∷SL2∷DsRed*]
TQ9467 *bbs-1(ok1111)* I; *xuEx1978*[*Psra-6∷GCaMP6f*+*Psra-6∷SL2∷DsRed*]
TQ9465 *bbs-2(ok2053)* IV; *xuEx1978*[*Psra-6∷GCaMP6f*+*Psra-6∷SL2∷DsRed*]
TQ9474 *arl-6/bbs-3(ok3472)* III; *xuEx1978*[*Psra-6∷GCaMP6f*+*Psra-6∷SL2∷DsRed*]
TQ9478 *bbs-4(tm3038)* III; *xuEx1978*[*Psra-6∷GCaMP6f*+*Psra-6∷SL2∷DsRed*]
TQ9475 *bbs-5(gk537)* III; *xuEx1978*[*Psra-6∷GCaMP6f*+*Psra-6∷SL2∷DsRed*]
TQ9476 *osm-12/bbs-7(ok1351)* III; *xuEx1978*[*Psra-6∷GCaMP6f*+*Psra-6∷SL2∷DsRed*]
TQ9468 *bbs-8(nx77)* V; *xuEx1978*[*Psra-6∷GCaMP6f*+*Psra-6∷SL2∷DsRed*]
TQ9473 bbs-9(gk471) I; *xuEx1978*[*Psra-6∷GCaMP6f*+*Psra-6∷SL2∷DsRed*]
TQ9636 *che-3(e1124)* I; *xuEx1978*[*Psra-6∷GCaMP6f*+*Psra-6∷SL2∷DsRed*]
TQ7538 *osm-3(p802)* IV; *xuEx1978*[*Psra-6∷GCaMP6f*+*Psra-6∷SL2∷DsRed*]
TQ9479 *kap-1(ok676)* II; *xuEx1978*[*Psra-6∷GCaMP6f*+*Psra-6∷SL2∷DsRed*]
TQ9634 *klp-11(tm324)* IV; *xuEx1978*[*Psra-6∷GCaMP6f*+*Psra-6∷SL2∷DsRed*]
TQ9737 *daf-10(e1387)* IV; *xuEx1978*[*Psra-6∷GCaMP6f*+*Psra-6∷SL2∷DsRed*]
TQ9639 *osm-5(p813)* X; *xuEx1978*[*Psra-6∷GCaMP6f*+*Psra-6∷SL2∷DsRed*]
TQ9749 *osm-6(p811)* V; *xuEx1978*[*Psra-6∷GCaMP6f*+*Psra-6∷SL2∷DsRed*]
TQ9629 *daf-19(m86)* II; *daf-12(sa204)* X; *xuEx1978*[*Psra-6∷GCaMP6f*+*Psra-6∷SL2∷DsRed*]
TQ9827 N2; *xuEx3318*[*Psra-6∷lite-1∷mNG∷3xFlag(cDNA)*+*Psra-6∷mCherry*]
TQ9890 N2; *xuEx3341*[*Psra-6∷lite-1∷mNG∷3xFlag(cDNA)*+*Psra-6∷mksr-2(cDNA)∷TagRFP*]
TQ7635 *osm-9(ok1677)* IV; *xuEx1978*[*Psra-6∷GCaMP6f*+*Psra-6∷SL2∷DsRed*]
TQ9702 *bbs-1(ok1111)* I; *lite-1*(*xu708*[LITE-1∷mNenoGreen∷3XFlag]) X
TQ9700 *bbs-8(nx77)* V; *lite-1*(*xu708*[LITE-1∷mNenoGreen∷3XFlag]) X
TQ9758 *bbs-1(ok1111)* I; *xuEx3272*[*Psra-6∷lite-1∷mNG∷3xflag(cDNA)*+*Punc-122∷rfp*]; *xuEx1978*[*Psra-6∷GCaMP6f*+*Psra-6∷SL2∷DsRed*]
TQ9715 *bbs-1(ok1111)* I; *xuEx3284*[*Psra-6∷bbs-1(cDNA)∷SL2∷mCherry*+*Punc-122∷gfp*]; *xuEx1978*[*Psra-6∷GCaMP6f*+*Psra-6∷SL2∷DsRed*]
TQ9717 *bbs-8(nx77)* V; *xuEx3286*[*Psra-6∷bbs-8(cDNA)∷SL2∷mCherry*+*Punc-122∷gfp*]; *xuEx1978*[*Psra-6∷GCaMP6f*+*Psra-6∷SL2∷DsRed*]
TQ9867 *bbs-8(nx77)* V; *xuEx3272*[*Psra-6∷lite-1∷mNG∷3xflag(cDNA)*+*Punc-122∷rfp*]; *xuEx1978*[*Psra-6∷GCaMP6f*+*Psra-6∷SL2∷DsRed*]
TQ9748 N2 *kyEx6191*[*Psra-9∷GCaMP5A*+*Pelt-2∷mCherry*]
TQ9618 *bbs-1(ok1111)* I; *kyEx6191*[*Psra-9∷GCaMP5A*+*Pelt-2∷mCherry*]
TQ9619 *bbs-8(nx77)* V; *kyEx6191*[*Psra-9∷GCaMP5A*+*Pelt-2∷mCherry*]
TQ7810 N2; xuIs471[*Pgpa-13∷FLP*ase+*Psra-6∷FRT2∷yfp2*]
TQ9746 *bbs-1(ok1111)* I; xuIs471[*Pgpa-13∷FLP*ase+*Psra-6∷FRT2∷yfp2*]
TQ9757 *bbs-8(nx77)* V; xuIs471[*Pgpa-13∷FLP*ase+*Psra-6∷FRT2∷yfp2*]

Worms were fed with standard *E. coli* strain OP50 on NGM agar plates at 20°C. Mutants were verified using PCR-based genotyping and sequencing. Worms carry extrachromosomal arrays were generated by injecting plasmid DNA into the hermaphrodite gonad. In all cases, the same extrachromosomal arrays were crossed into wild-type N2 and different mutant backgrounds for expression analysis and calcium imaging.

### Genetic screen

*bbs-1(xu583), osm-12(xu563), bbs-8(xu607), bbs-8(xu608)*, and *bbs-9(xu579)* were isolated from a forward genetic screen for mutants with light insensitivity phenotype in ASH neurons. *xuIs556* worms co-expressing *Psra-6∷case12* and *Psra-6∷SL2∷mCherry* were mutagenized with 50 mM EMS. 30-50 F2 worms from each F1 plate (2 F1 per plate) were checked directly under a fluorescence dissecting microscope (Zeiss Discovery) coupled with a SOLA light engine LED light source (Lumencor). F2 worms were examined with 5 mW/mm^2^ blue light under 1x lens. Light intensity was measured using an optometer (S471, UDT Instruments) coupled with a sensor head (268LP, UDT Instruments). A total of ~36,000 F2 animals were screened. Animals exhibiting no light response, but with normal-looking soma and dendrites in ASH neurons, were picked, and their progeny was re-examined to confirm the phenotype. Counter-screens were performed using 1 M glycerol. Animals exhibiting normal glycerol responses were kept for further analysis.

Mutant strains were crossed with Hawaii strain CB4856, F2 homozygous mutant worms were pooled, and their genomic DNA was extracted and purified using a NucleoSpin DNA RapidLyse kit (Cat. No. 740100) from MACHERY-NAGEL and subjected for pair-end Illumina whole-genome sequencing ^42^. Unique variants were identified using the MiModD software package via the Variant Allele Frequency (VAF) mapping method (https://mimodd.readthedocs.io/en/latest/#). Protein coding variants were identified using SnpEff version 4.3 (http://snpeff.sourceforge.net) and the WS235 reference genome (http://wormbase.org).

### Molecular biology

cDNA of *bbs-1, bbs-8,* and *mksr-2* was amplified by RT-PCR from N2 worms and inserted downstream of the *sra-6* promoter. *lite-1∷mNeonGreen* cDNA was amplified from *xu708* CRISPR/Cas9 knock-in strain by RT-PCR. To visualize the transition zone and basal bodies of ASH neurons, cDNA of *mksr-2* was fused in frame with a TagRFP reporter gene at the N-terminus ^43^.

### Generation of *lite-1(xu492)* and *lite-1(xu708)* alleles

*lite-1(xu492)* is a deletion allele with a 2701 bp deletion generated by CRISPR/Cas9-based gene editing using a protocol developed by the Fire lab ^44^. Specifically, four sgRNAs targeting exon 1 (5’-TTTGAATGTGATGATGGTGG-3’, 5’-ATGTTTGAATGTGATGATGG-3’), exon 4 (5’-ACTTTTGGCTCTTACCATGG-3’), and exon 6 (5’-GTAGAACAAGATTGCCAAGG-3’) were cloned into the pU6 plasmid and injected together with *Peft3∷Cas9* and *dpy-10* sgRNA plasmids, and ssDNA donor for *dpy-10(cn64)*. Both roller and dumpy F1 worms were singled onto NGM plates, and animals with deletions were selected by single worm PCR. The deletion was verified by Sanger sequencing with and the flanking sequences are: 5’-CGTAAAAAACAACATGCCACCAC-2701bp deletion - GGCGGCCACCTACGCCAGTA -3’.

*lite-1(xu708)* is a knock-in allele with an mNeonGreen∷3xFlag tag attached to the C-terminus of *lite-1* locus by the self-excising cassette (SEC) method developed by Goldstein lab ^45^. Four sgRNAs targeting the last exon and 3’-UTR region (5’-TTGCGATATTCTGGAGACTC-3’, 5’-TCGTGTGGTTTGCGATATTC-3’, 5’-GCTTTTATGTGTGAATCGTG-3’, and 5’-AGTGACAGCTGAAGATAAAA-3’) near the stop codon were cloned into the pU6 plasmid and injected together with Peft3∷Cas9, SEC vector pDD268 with 500-700bp homology arms, and Pmyo-3∷mCherry marker. Homozygous rollers were selected using hygromycin, and the selectable marker was removed following heat shock. The insertion was confirmed by sequencing.

### Microscopy and calcium imaging

Worms were mounted on 2% agarose pads with 20 mM sodium azide (Sigma-Aldrich, Cat. No. S2002) or tetramisole (Sigma-Aldrich, Cat. No. T1512) in M9 solution. Images were acquired with a Nikon A1 confocal microscope with a 60X lens and were processed using ImageJ.

Day 1 adult worms were used in all experiments unless noted otherwise. Animals were immobilized using cyanoacrylate adhesive glue to record light response or using microfluidic chips to record chemical responses. All GCaMP signals were recorded with MetaFluor software (Molecular Devices) and a Hamamatsu sCMOS camera (Orca Flash 4.0 LT) using a 40x lens on Olympus IX73 microscope. Worms carrying a GCaMP6f transgene in ASH neurons and GCaMP5A in ASK neurons were used for calcium imaging. 20 μL M9 buffer was added to cover the worm glued on a 2% agarose pad. Worms were allowed to rest for 3 min before 5.6 mW/mm2 480 nm blue light stimulus was applied to excite GCaMP and trigger the photoresponse in ASH. Images were acquired at 10 Hz frame rate with 100 ms exposure time. A total of 700 frames were recorded for each worm.

To record ASH response to 1 M glycerol, 1 mM CuCl2, and 0.1% SDS, worms were immobilized in a microfluidic chip as previously described ^30,46^. All chemicals were dissolved in M13 buffer (30 mM Tris-HCl, 100 mM NaCl, 10 mM KCl, pH 7.0). Worms were imaged with 480 nm and 565 nm light spontaneously, and both GCaMP fluorescence and dsRed fluorescence were recorded. Worms were pre-exposed to imaging light for 1-5 min to quench the photoresponse in ASH neurons. After the basal line became stable, a 30 s stimulus pulse was applied to the worm. Each worm was challenged once only with the stimulus. Data were presented as ΔR/R0 (ΔR=R−R0, R=F_GCaMP6f_/F_dsRed_).

### Phototaxis behavior

Phototaxis behavior was analyzed as described previously ^26^. Briefly, day 1 adult worms were transferred to NGM plates covered with a thin layer of freshly spread OP50 bacteria ~30 min before the test. Worms were tested under a 10x objective on a fluorescence dissection microscope (Zeiss Discovery). A 2 s light pulse (350±25 nm, 406 μW/mm2, Sola, Lumencor) was delivered to the head of a worm that was slowly moving forward. A positive response was scored if the worm stopped forward movement during the illumination time or 3 s after and initiated a backward movement that lasted at least half a head swing.

### qRT-PCR analysis

ASH neurons were marked by *xuIs471*[*Pgpa-13∷FLPase*+*Psra-6∷FRT2∷yfp2*] transgene. ASH neurons were isolated from wild-type, *bbs-1(ok1111)* or *bbs-8(nx77)* adult worms carrying *xuIs471* transgene by fluorescence flow cytometry (FACSAria III cell sorter) using a protocol described previously ^36^. TRIzol LS (Invitrogen, Cat. No. 10296010) was used to extract the total RNA from ASH neurons. RNA samples were further treated with DNaseI (Qiagen, Cat. No. 79254) and cleaned using RNeasy MinElute Cleanup Kit (Qiagen, Cat. No. 74204). High-Capacity cDNA Reverse Transcription Kit (Applied Biosystems, Cat. No. 4368814) was used to synthesize cDNA using oligo dT primer (Invitrogen, Cat. No. 18418012). SYBR Green was used for qPCR analysis (Applied Biosystems, Cat. No. A25742), and the samples were run on a QuantStudio™ 5 System (Applied Biosystems, Cat. No. A33628) using a 384-well plate. *act-1* and *pmp-3* were used as housekeeping genes for *lite-1* gene expression analysis. *act-1* forward primer is 5’-CCAGGAATTGCTGATCGTATGCAGAA-3’, and the reverse primer is 5’-TGGAGAGGGAAGCGAGGATAGA-3’. *pmp-3* forward primer is 5’-TGGCCGGATGATGGTGTCGC-3’, and the reverse primer is 5’-ACGAACAATGCCAAAGGCCAGC-3’. *lite-1* forward primer is 5’-TGCTGGCTTCATTGCAACTAT-3’, and the reverse primer is 5’-TTCAACGAAAACTGGCACAA-3’. The experiments were designed to include three biological repeats and three technical replicates for each target.

### Statistics and Data Analysis

Statistical analysis was performed in GraphPad Prism.

## Acknowledgments

We thank the Caenorhabditis Genetics Center (CGC), National BioResource Project (NBRP), and Cori Bargmann for strains, and Patrick McGrath for CRISPR/Cas9 plasmids. This work was supported by the NIGMS (to X.Z.S.X.).

---

**Figure S1.**
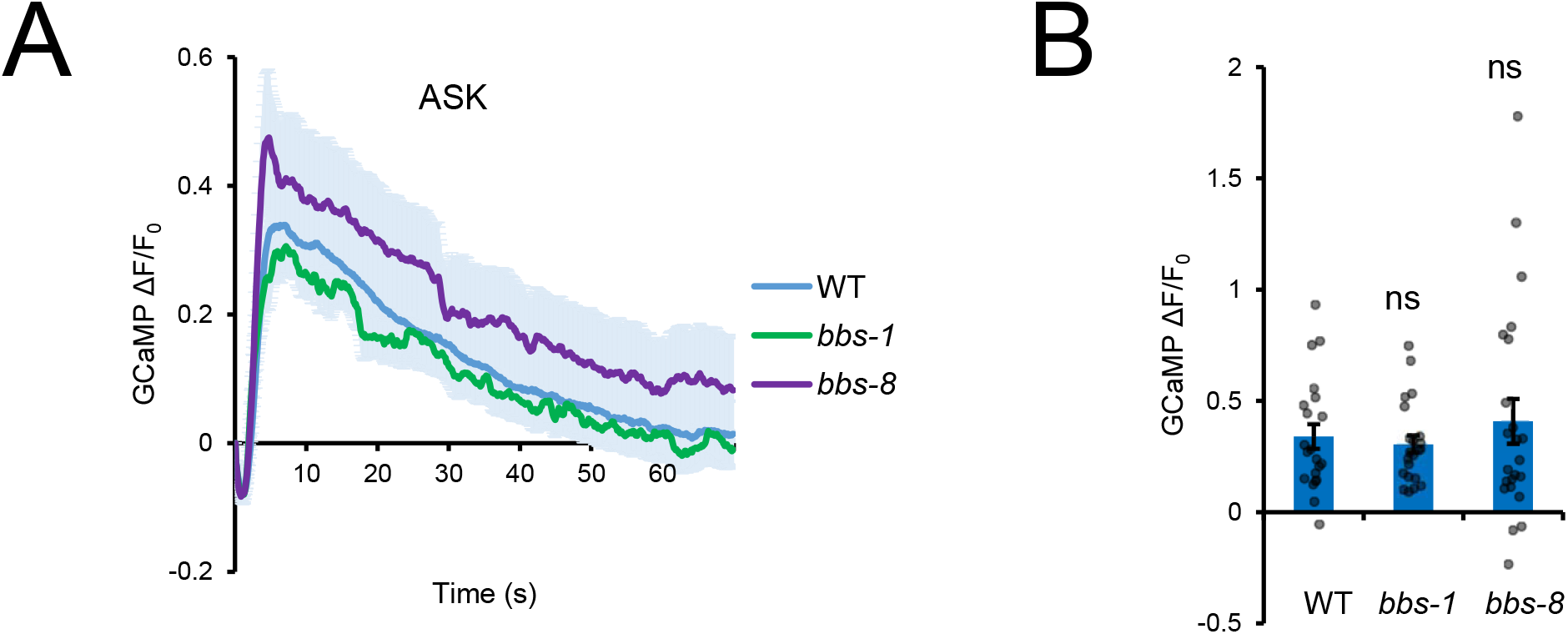
BBSome is not required for ASK neurons to sense light. ASK neurons respond normally to light in *bbs-1(ok1111)* and *bbs-8(nx77)* mutant worms. All worms carried the same GCaMP5A transgene expressed in ASK neurons. **(A)** Calcium imaging traces. Shades along the traces denote error bars (SEM). **(B)** Bar graph. Statistics were calculated using One-way ANOVA Bonferroni test, and all strains were compared to wild type. n≥21. ns: not significant.ificant.

**Figure S2.**
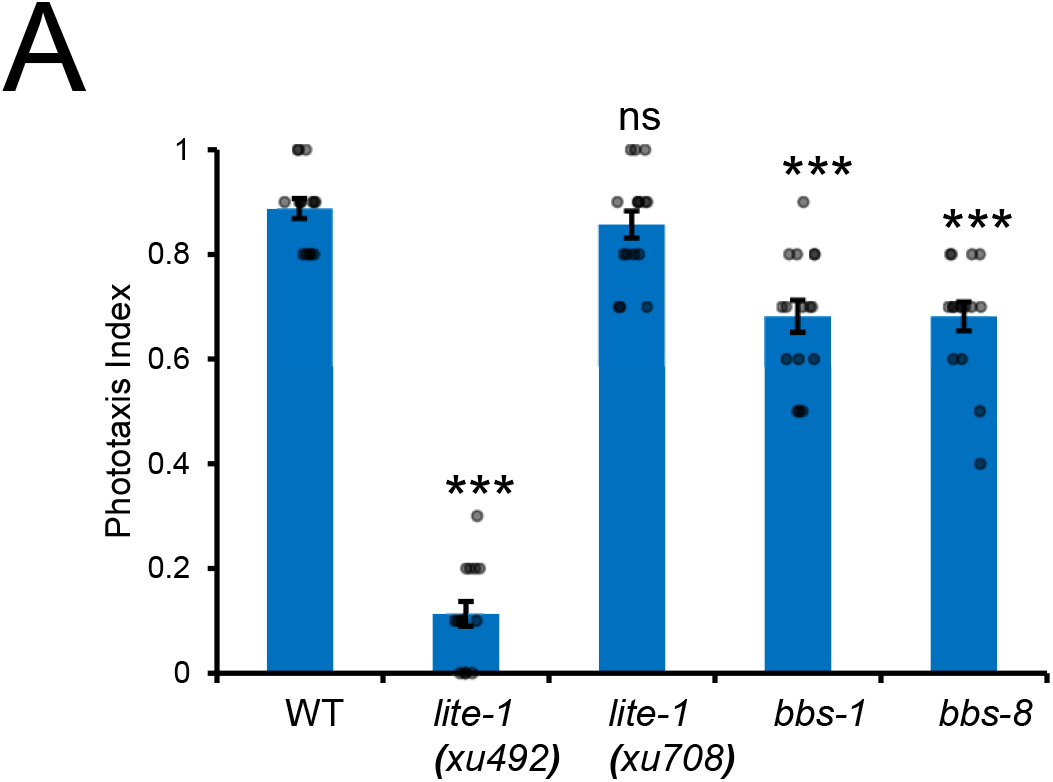
Phototaxis behavior in *bbs* mutants and *lite-1* knockin and knockout alleles. Phototaxis assay shows that *bbs-1(ok1111)* and *bbs-8(nx77)* mutants have modest defects compared to wild type. This is expected, as BBSome only affects the light sensitivity of ASH neurons but not other photosensory neurons. While the knock-out null allele *lite-1(xu492)* was severely defective in phototaxis, the knock-in allele *lite-1(xu708[LITE-1∷mNeonGreen])* was normal, indicating that the nNeonGreen tag fused to the C-terminus of LITE-1 does not affect LITE-1 function. n≥15. Error bars: SEM. ***P<0.0001 (one-way ANOVA with Bonferroni test).

**Figure S3.**
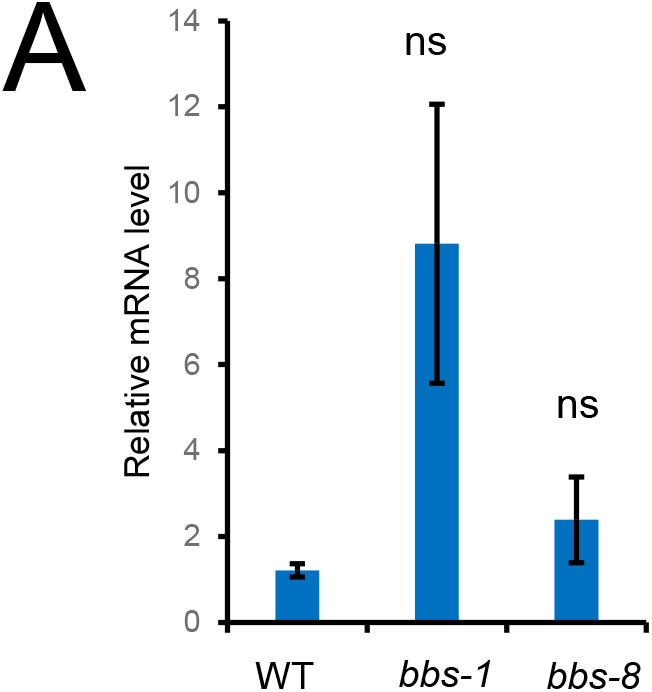
*Lite-1* mRNA level in ASH neurons is normal in *bbs* mutant worms. qRT-PCR results show that the *lite-1* mRNA level in ASH neurons is similar between wild type, *bbs-1(ok1111)* and *bbs-8(nx77)* strains. ASH neurons were isolated by fluorescence flow cytometry, and their total RNA were extracted and subjected to qRT-PCR analysis. Shown are data from three replicates, and the experiment was repeated three times. Error bars: SEM. Statistics were calculated using One-way ANOVA Bonferroni test, and all strains were compared to wild type. ns: not significant.

